# Attenuating ABHD17 isoforms augments the *S*-acylation and function of NOD2 and a subset of Crohn’s disease-associated NOD2 variants

**DOI:** 10.1101/2023.12.20.572362

**Authors:** Charneal L. Dixon, Noah R. Martin, Micah J. Niphakis, Benjamin F. Cravatt, Gregory D. Fairn

## Abstract

**BACKGROUND AND AIMS:** NOD2 is an intracellular innate immune receptor that detects bacterial peptidoglycan fragments. Although nominally soluble, some NOD2 is associated with the plasma membrane and endosomal compartments for microbial surveillance. This membrane targeting is achieved through post-translational *S*- acylation of NOD2 by the protein acyltransferase ZDHHC5. Membrane attachment is necessary to initiate a signaling cascade in response to cytosolic peptidoglycan fragments. Ultimately, this signaling results in the production of antimicrobial peptides and proinflammatory cytokines. In most cases, *S*-acylation is a reversible post-translational modification with removal of the fatty acyl chain catalyzed by one of several acyl protein thioesterases. Deacylation of NOD2 by such an enzyme will displace it from the plasma membrane and endosomes, thus preventing signaling.

**METHODS:** To identify the enzymes responsible for NOD2 deacylation, we used engineered cell lines with RNA interference and small-molecule inhibitors. These approaches were combined with confocal microscopy, acyl-resin-assisted capture, immunoblotting, and cytokine multiplex assays.

**RESULTS:** We identified α/β-hydrolase domain-containing protein 17 isoforms (ABHD17A, ABHD17B, and ABHD17C) as the acyl protein thioesterases responsible for NOD2 deacylation. Inhibiting ABHD17 increased the plasma membrane localization of wild-type NOD2 and a subset of poorly acylated Crohn’s disease-associated variants.

This enhanced NOD2 activity, increasing NF-κB activation and pro-inflammatory cytokine production in epithelial cells.

**CONCLUSIONS:** These findings demonstrate that ABHD17 isoforms are negative regulators of NOD2. The results also suggest that targeting ABHD17 isoforms could restore functionality to specific Crohn’s disease-associated NOD2 variants, offering a potential therapeutic strategy.

**Grant Support:** This work was supported by a Project Grant from the Canadian Institutes of Health Research (grant no.: PJT166010; to G.D.F.), an Innovator Award from the Kenneth Rainin Foundation, and a grant from the National Institutes of Health, (R01CA193994 to B.F.C). A Tier 1 Canada Research Chair supports G.D.F. in Multiomics of Lipids and Innate Immunity. C.L.D. was supported by a Breakthrough Accelerator Fellowship from the Dalhousie Medical Research Foundation/Medical Research Development Office. N.M is a recipient of a graduate scholarship from the I3V Wave and the Dalhousie Medical Research Development Office.

**Disclosures:** The authors declare that the research was conducted in the absence of any commercial or financial relationships that could be construed as a potential conflict of interest.

**SYNOPSIS:** The peptidoglycan sensor NOD2 requires post-translational *S*-acylation to associate with cellular membranes and transduce signals. This study identified the ABHD17 family of thioesterases as responsible for NOD2 deacylation and inactivation. Inhibiting or silencing ABHD17 isoforms increases *S*-acylation and functionality of NOD2 and a subset of Crohn’s disease-associated variants.

## INTRODUCTION

Crohn’s disease (CD) is a prominent form of inflammatory bowel disease that can affect any segment of the gastrointestinal tract, extending from the mouth to the anus, that most commonly manifests in the lower portion of the small intestine and the upper colon. CD is characterized by a range of symptoms, including weight loss, fatigue, diarrhea, and abdominal pain, which can be debilitating and, in some instances, life-threatening. Current treatments, including the use of steroids and tailored dietary approaches, have proven effective in alleviating these symptoms and forestalling disease exacerbations. Despite these advances, CD remains an intricately multifactorial condition influenced by genetic predisposition, environmental factors, dietary habits, and an individual’s unique microbiome.

The nucleotide-binding oligomerization domain-containing protein 2 (NOD2) gene has been consistently implicated in the pathogenesis of CD since it was first identified in 2001 [1–3]. NOD2 is a member of the evolutionarily conserved family of NOD-like receptors (NLR) that provide intracellular surveillance through detecting cytoplasmic pathogen-associated molecular patterns [4–6]. NOD2 belongs to the NLRC subfamily, containing two amino-terminal caspase activation and recruitment domains (CARD) [7]. NOD2, and the closely related NOD1, bind small peptides originating from bacterial peptidoglycan. The binding of ATP and peptidoglycan fragments to NOD2 induces conformational changes and oligomerization, engaging receptor-interacting protein kinase 2 (RIPK2) and triggering a cascade of ubiquitination and phosphorylation events. This process activates the nuclear factor kappa B (NF-lrlB) and mitogen-activated protein kinases (MAPK) pathways, ultimately inducing the transcription of pro-inflammatory cytokines, chemokines, and other inflammatory mediators [8, 9].

The complex pathophysiology of CD has led to the emergence of multiple functional roles for NOD2. In the presence of a diverse and healthy microbiome, NOD2 is thought to provide signals that facilitate the expression and secretion of small anti- microbial peptides (AMPs) such as lil- and β-defensins by Paneth cells and other epithelial cells of the small and large intestine [10, 11]. Thus, loss of NOD2 function and a reduction in AMPs can lead to changes in the microbiome and dysbiosis. This shift in the bacterial population and activity can result in an increase in bacteria capable of damaging or invading the epithelial lining. In this instance, NOD2, in combination with ATG16L1, another CD risk factor, is critical in initiating an anti-bacterial autophagic pathway in response to invasion [12–14]. Finally, NOD2 can be exposed to peptidoglycan in resident macrophage and epithelial cells through its uptake via the SLC15A family of transporters in the plasma membrane (PM) and endosomes to initiate inflammatory cytokine production [15–17]. However, precisely how loss of NOD2 contributes to the development of CD remains unclear, although it may impair the ability to recover from acute infections, as observed in murine models [18].

In contrast to toll-like receptors that recognize extracellular or luminal bacterial components, NOD2 is an intracellular receptor that attaches to the surface of the PM and endomembranes to transduce signals following binding to phosphorylated muramyl dipeptide (MDP) in the cytosol [17, 19]. Previously it was demonstrated that *S-*acylation (often called *S*-palmitoylation or palmitoylation), was required to target NOD2 and NOD1 to membranes [20, 21]. It was further revealed that the protein acyltransferase ZDHHC5 mediates the attachment of the fatty acyl group to NOD2 (Cys395 and 1033) and NOD1 (Cys558, 567, 952); this modification is critical for the ability of NOD proteins to respond to peptidoglycan components and mount an effective inflammatory response [21].

Additionally, several CD-associated loss-of-function variants of NOD2 exhibit reduced levels of *S*-acylation, referred to as hypo-acylation. The *S-*acylation deficiency correlates with decreased signaling in response to peptidoglycan for these hypo-acylated variants. As *S-*acylation is a reversible post-translational modification, we theorized that identifying and inhibiting the responsible acyl protein thioesterase(s) could potentially enhance the *S*-acylation of not only wild-type NOD2 but also hypo-acylated mutants.

Indeed, such an intervention could improve the functionality of some of the CD-associated NOD2 variants.

In this study, we identify the family of lrl/lrl-hydrolase domain-containing protein 17 (ABHD17A, 17B, 17C) as the enzymes responsible for the deacylation of NOD2.

Furthermore, we find that inhibiting ABHD17 enzymes partly restores the localization, *S*- acylation, and ligand-induced signaling of a subset of CD-associated loss-of-function NOD2 variants.

## RESULTS

### Deacylation of NOD2 is Enzymatically Mediated

Unlike other types of protein lipidation, the covalent attachment of palmitate and other fatty acids to cysteine residues via a thioester bond is reversible [22]. Dynamic changes in *S-*acylation status play a role in the regulation and function of proteins [23–25]. Indeed, the proportion of NOD2 that are *S-*acylated is increased in response to stimuli [21]. Yet, to date, nothing is known about the reversibility of this modification on NOD2 or the identity of an acyl protein thioesterase (deacylating) enzyme.

To determine if colorectal epithelial cells express a thioesterase capable of deacylating NOD2, we first developed a cell model with controllable low-level expression of GFP-NOD2. To do this, we used the Sleeping Beauty transposon system to engineer HCT116 and HT29 epithelial cell lines with a doxycycline-inducible GFP-tagged NOD2 (GFP- NOD2) as described in the Methods. Following the generation and selection of the polyclonal cell line, doxycycline-inducible expression of GFP-NOD2 was confirmed by immunoblotting and microscopy (Figure 1A, *B*). In these cells, there was a sizeable cytosolic pool of GFP-NOD2 and a smaller fraction associated with the PM, consistent with previous results [21, 26, 27]. Palmostatin B is a cell-permeable, broad-spectrum inhibitor of acyl protein thioesterases, including acyl protein thioesterase (APT) 1 and 2 [28, 29], palmitoyl-protein thioesterase 1 [29], α/β-hydrolase domain (ABHD) 6 [30], ABHD10 [31] ABHD12 [32], ABHD16A [30, 33] and ABHD17(A/B/C) [30].

**Figure 1.**
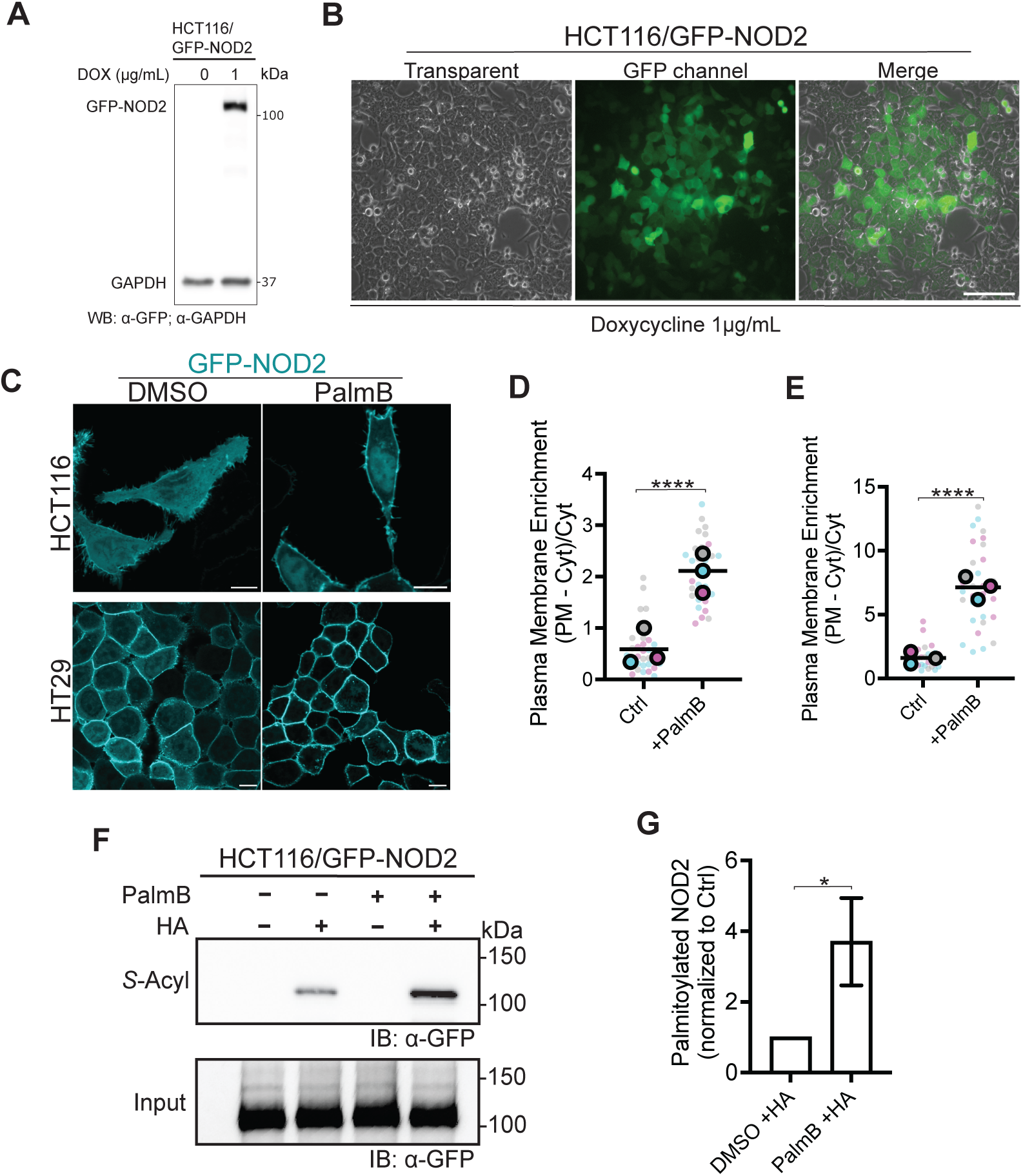
Inhibiting acyl protein thioesterases increases plasmalemmal NOD2 and *S*-acylation. (*A*) Immunoblot showing doxycycline-inducible expression of GFP-NOD2 and GAPDH in HCT116 cells. (*B*) Low magnification epifluorescence micrographs of GFP-NOD2 in HCT116 cells following induction with 1 μg/mL doxycycline for 18 h. Scale bar, 100 μm. (*C*) Confocal microscopy of human colorectal carcinoma cells HCT116 and HT29 expressing stably transfected GFP-NOD2 (pseudo-colored cyan). Cells were treated with 100 μM palmostatin B (PalmB) or dimethyl sulfoxide (DMSO, vehicle) for 4 h. Scale bars, 10μm. (*D*) Quantification of plasma membrane localization and the effect of PalmB on GFP-NOD2 in HCT116 (*D*) and HT29 (*E*) cells from images as in (*C*). (*F*) *S*-acylation levels of GFP-NOD2 expressed in HCT116 cells treated with DMSO or PalmB. *S*-acylated GFP-NOD2 was detected in total cell lysates using acyl resin-assisted-capture (acyl-RAC) assay and immunoblot analysis; samples omitting hydroxylamine (HA) are used as a control to demonstrate the fidelity of the assay specificity. (*G*) Quantification of *S*-acylation assay depicted in (*F*). (*H*) Average fold change quantification for quantification shown in (G). All data are representative of at least three independent experiments, with means indicated by a horizontal black line and samples from individual replicates color-matched. Unpaired two-tailed Student’s t-test determined P-values for data in D, E, and G; p≤0.05 (*) and p≤0.0001 (****).

Thus, we next treated cells expressing GFP-NOD2 with either solvent control (DMSO) or Palmostatin B for 4 h. Following the 4 h incubation with Palmostatin B, confocal microscopy revealed a significant enrichment of the GFP-NOD2 on the PM (p<0.0001) (*Figure 1C-E)*.

Since it was previously demonstrated that the NOD2^C395,1033S^ S-acylation deficient mutant is completely cytosolic [20, 21], we reasoned that the observed plasmalemmal enrichment was due to a more substantial fraction of GFP-NOD2 being *S-*acylated. To confirm that Palmostatin B increases S-acylation of NOD2 experimentally, we used an acyl resin-assisted capture (acyl-RAC) technique (*Figure 1F*, G) that uses a series of steps, including the blockage of free-thiols, cleavage of acyl-thioester bonds, and the use of an affinity matrix to bind the newly liberated and previously *S*-acylated cysteine residues from cell lysates. Immunoblotting GFP-NOD2 following acyl-RAC demonstrated a 3.2-fold increase (p=0.0192) in *S-*acylated GFP- NOD2 in HCT116 cells treated with Palmostatin B compared to control (*Figure 1H*). The presence of a signal with hydroxylamine (+HA) treatment indicates that fatty acids are attached to the protein via a thioester bond. These results suggest that *S-*acylation of NOD2 is reversible and that a Palmostatin B-sensitive protein mediates the enzymatic removal of acyl chains from NOD2.

### Inhibiting ABHD17 Enzymes Increases NOD2 S-acylation

Motivated by the impact of Palmostatin B on NOD2 localization and *S-*acylation, we used selective small-molecule inhibitors to target known deacylation enzymes to identify the relevant NOD2 thioesterases. HCT116 cells stably expressing GFP-NOD2 were treated with DMSO or inhibitor for a total of 4 h. Treatment with ML348 [34] and ML349 [34], which target APT1 and APT2, respectively, had no discernable effect on the localization of GFP-NOD2 (*Figure 2A*). A similar lack of response was observed with the selective ABHD6 inhibitor, WWL70 [35, 36]. Conversely, THL [36, 37] and ABD957 [38] treated HCT116 cells have significant enrichment of GFP-NOD2 at the PM, 2.2-fold (p=0.0175) and 4.9-fold (p<0.0001) increase, respectively (*Figure 2B*). A similar enrichment was observed in HT29 cells (p<0.0001) (*Figure 2C*, *D*). For ABD957-treated HCT116 cells, this corresponded to a mean fold increase of 4.4 (p=0.0061) in the fraction of *S*-acylated NOD2 compared to DMSO, as determined by the acyl-RAC assay. In contrast, the THL treatment did not significantly impact the amount of *S-* acylated NOD2 protein captured during the assay (p=0.3455) (*Figure 2E, F*).

**Figure 2.**
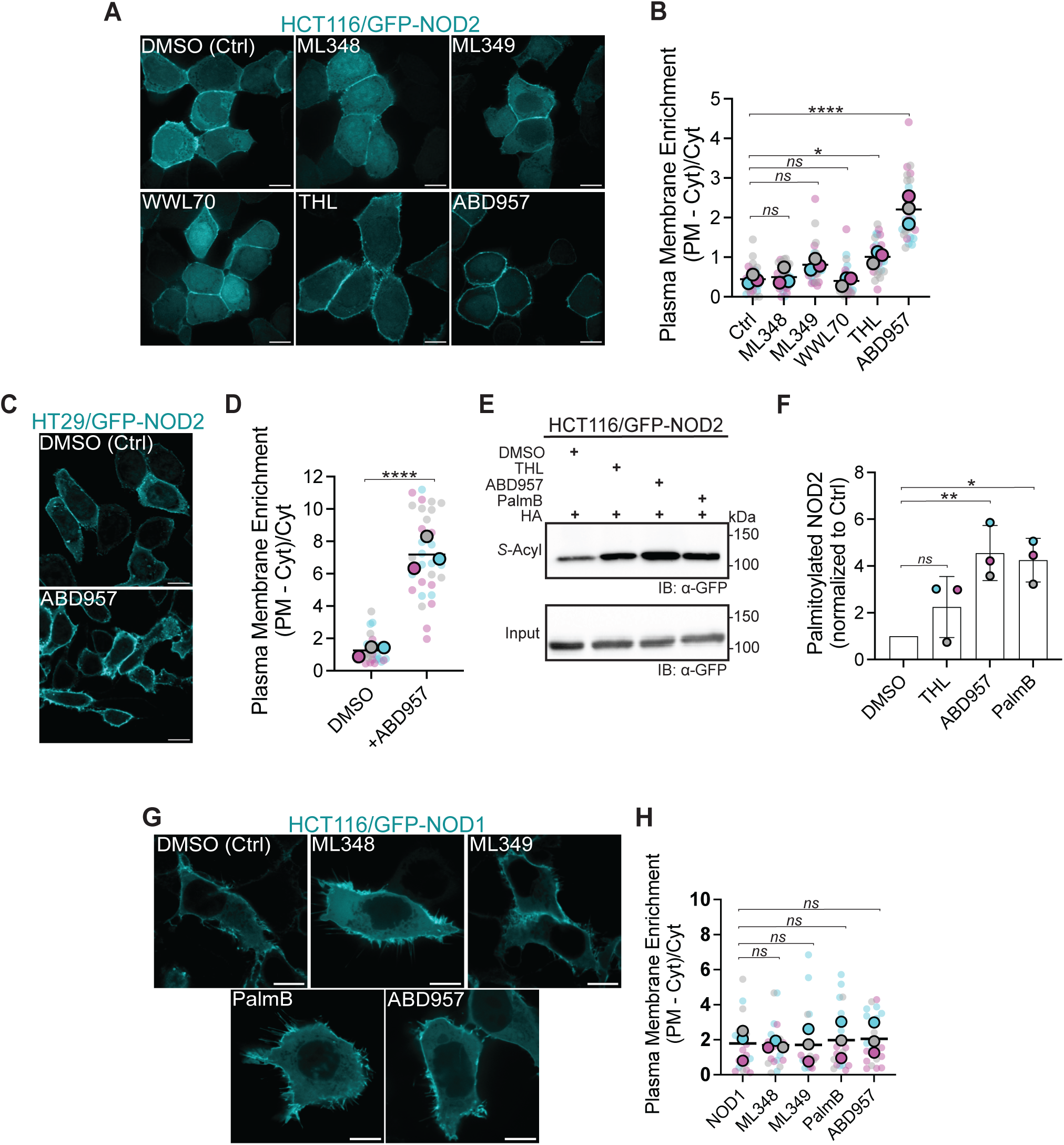
ABHD17 isoforms mediate enzymatic deacylation of NOD2. (*A, G*) Confocal images showing localization of GFP-NOD2 (A) and GFP-NOD1 (G) in HCT116 cells treated with selective small molecule acyl thioesterase inhibitors: 2.5 μM ABD957, 50 μM ML348 or 40 μM ML349 for 4 h. GFP-NOD2 expressing cells were also treated with 40 μM orlistatin (THL) and 25 μM WWL70. Scale bars, 10μm. (*B, H*) Quantification of GFP-NOD2 and GFP-NOD1 plasma membrane localization from images as in (*A, G*). (*C*) Confocal images showing localization of GFP-NOD2 in HT29 cells treated with 2.5 μM ABD957. (*D*) Quantification of GFP-NOD2 plasma membrane localization from representative images as in (C) in HT29 cells treated with ABD957 or DMSO. (*E, F*) Acyl-RAC analysis of GFP-NOD2 in HCT116 cells treated with THL, ABD957 or PalmB. All data are representative of at least three independent experiments, with means indicated by a horizontal black line and samples from individual replicates color-matched. Statistical significance was determined by two-tailed Student’s unpaired *t*-test or one-way ANOVA followed by Dunnett’s *post hoc* test. Group differences were considered significant for p≤0.05 (*), p≤0.01 (**) and p≤0.0001 (****). Not significant; *ns*.

These data suggest that at least one of the ABHD17 isoforms may be the primary enzyme responsible for deacylating NOD2. We further hypothesized that given the homology between NOD1 and NOD2 and that ZDHHC5 is required for the *S*-acylation of both NOD1 and NOD2, the same deacylating enzyme may regulate NOD1. To test this, we treated HCT116 cells stably transfected with doxycycline-inducible GFP-NOD1 with the various thioesterase inhibitors but observed no significant increase (p>0.05) in plasma membrane enrichment (*Figure 2G*, *H*). However, given the limited cytoplasmic pool of GFP-NOD1, any observed change in plasma membrane enrichment could be negligible due to the minimal amount of soluble NOD1 available for *S*-acylation.

### Overexpression of ABHD17 Enzymes Depletes the Plasmalemmal Pool of NOD1 and NOD2

Having narrowed the list of candidate enzymes using inhibitors, we next examined the cellular consequences of expressing the individual α/β-hydrolases targeted by THL and ABD957. Preventing NOD2 *S-*acylation prevents NOD2 and NOD1 from localizing to the PM [20, 21]. Accordingly, we anticipated that overexpressing an α/β-hydrolase domain enzyme capable of deacylating NOD2 would similarly reduce the fraction of GFP-NOD2 associated with the PM. HCT116 cells stably expressing GFP- NOD2 were transiently transfected with individual ABHD enzymes (6, 12, 16A,17A, 17B and 17C) (*Figure 3A*). The three ABHD17 isoforms significantly redistributed GFP- NOD2 from PM to cytosol, with ABHD17B and 17C being the most effective in this assay (*Figure 3B*). In contrast, overexpression of the THL-sensitive ABHD6, ABHD12, and ABHD16A did not significantly deplete the plasmalemmal GFP-NOD2 signal (p>0.05). These data further support ABHD17 isoforms as the primary NOD2 deacylation enzymes. When we applied this strategy to HCT116 cells stably expressing GFP-NOD1 (*Figure 3C*, *D*), we observed a modest ≈ 40%reduction (p=0.009) in GFP- NOD1 signal at the PM when ABHD17A was overexpressed. ABHD17B and 17C isoforms were more effective (p<0.0001) at redistributing GFP-NOD1 from the PM to cytosol, with ≍ 60% and ≍ 78% decrease, respectively. These results support the notion that ABHD17 isoforms are the relevant deacylating enzymes and, thus, negative regulators of NOD1 and NOD2 function.

**Figure 3.**
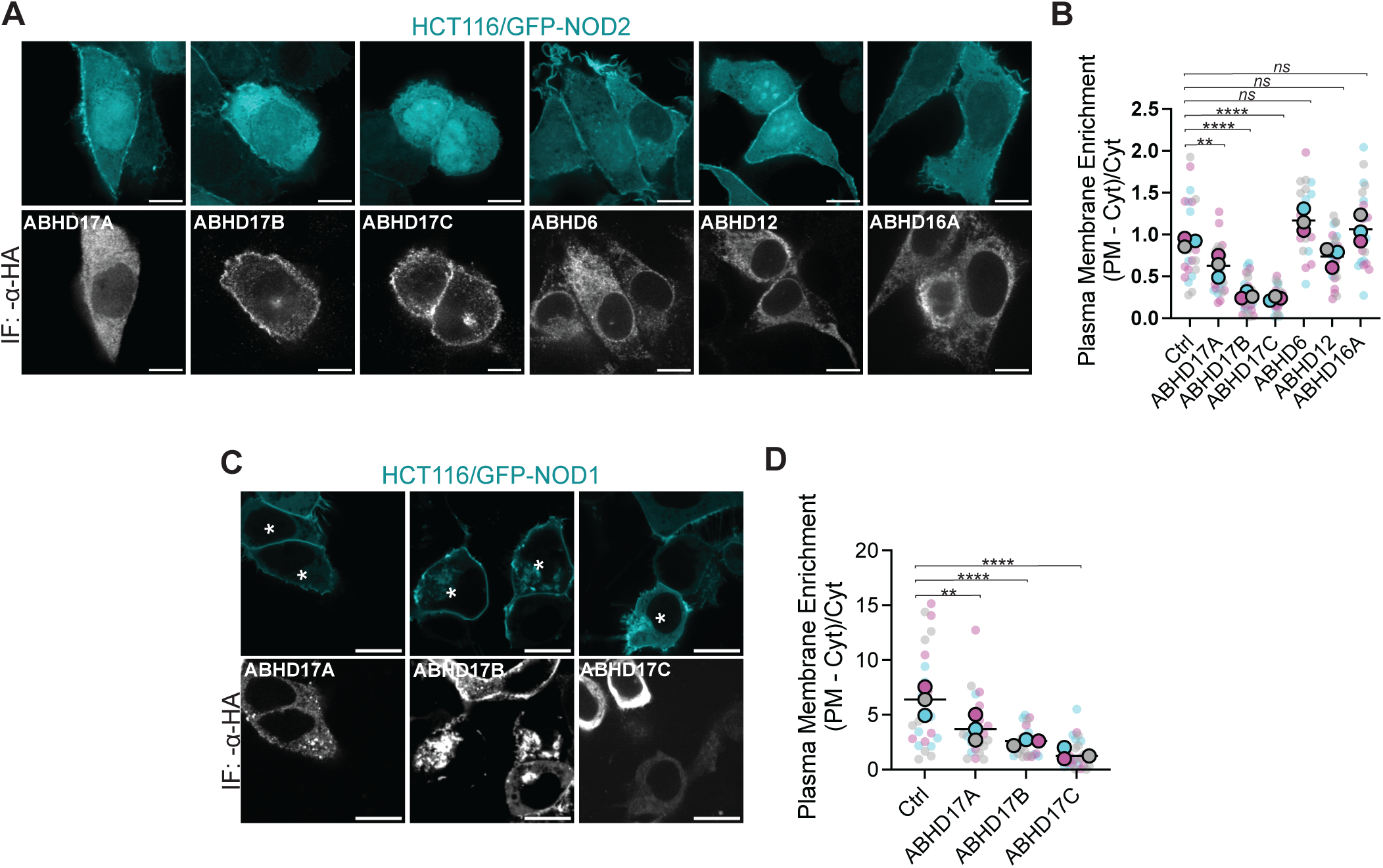
Enforced expression of ABHD17 decreases NOD1 and NOD2 plasma membrane localization. (*A*) Confocal images of stable HCT116/GFP-NOD2 (cyan) cells transiently transfected with 3HA-tagged a//3-hydrolase domain (ABHD) 6, 12, 16A, 17A, 17B and 17C enzymes (greyscale). (*B*) Quantification of GFP-NOD2 plasma membrane localization from images as in *A*. (*C*) Confocal images of stable HCT116/NOD1 (cyan) transiently transfected with 3HA-tagged ABHD17 isoforms (greyscale). Asterisks (*) indicate GFP-NOD1 and ABHD17 co-transfected cells. All data are representative of at least three independent experiments, with means indicated by a horizontal black line and samples from individual replicates color-matched. P-values for B-D and F-H were determined by ordinary one-way ANOVA followed by Dunnett’s multiple comparison test.; p≤0.01 (**), p≤0.0001 (****). Not significant; *ns*. Scale bars, 10 μm.

### Attenuating ABHD17 Increases the Plasmalemmal Enrichment of NOD2

ABHD17 enzymes are ubiquitously expressed in most cell types (Figure *S1A, B*), while NOD2 expression is restricted to immune cells and intestinal epithelial and Paneth cells [39, 40]. Although this suggests a broader regulatory role for these AHBD17 enzymes, the high expression of ABHD17C in gastrointestinal tissues makes it a key candidate for influencing NOD2 function. The ubiquitous expression of ABHD17 isoforms, particularly the elevated levels of ABHD17C in the gastrointestinal tract, supports its relevance to gastrointestinal immunity. Next, we investigated the effects of silencing the individual ABHD17 on NOD2 *S-*acylation in HCT116 cells. The relative expression of ABHD17 isoforms was analyzed using quantitative polymerase chain reaction (qPCR). All three isoforms have similar mRNA levels in HCT116 cells (*Figure 4A*). Thus, we conducted experiments by knocking down individual isoforms and all three simultaneously. Reverse transcription (RT)-qPCR showed efficient silencing of ABHD17A, 17B, and 17C either individually or in combination 48 hours after introduction of siRNA (Figure 4B). Silencing of ABHD17A had no significant effect on NOD2 localization (p= 0.1306). However, this siRNA only achieved ∼75% reduction in the transcript (Figure 4C, D), raising the possibility that the lack of effect is due to incomplete silencing. Conversely, knockdown of either ABHD17B or ABHD17C moderately affected GFP-NOD2 membrane localization, resulting in a 1.8-fold (p=0.0007) and 1.6-fold (p=0.0203) increase, respectively. Notably, the simultaneous knockdown of all three ABHD17 isoforms resulted in a 3.9-fold increase (p<0.0001) in GFP-NOD2 associated with the plasma membrane. The increase in the triple knockdown cells (Triple KD) was comparable to cells treated with the selective ABHD17 inhibitor (ABD957) (Figure 2B). The results from overexpression and silencing experiments indicate that the ABHD17 isoforms, particularly ABHD17B and ABHD17C, are the main thioesterases responsible for NOD2 deacylation in HCT116 cells. These findings are consistent with ABHD17 expression data, which show that ABHD17A, ABHD17B, and ABHD17C are expressed at levels higher than NOD2 (Figure S1).

**Figure 4.**
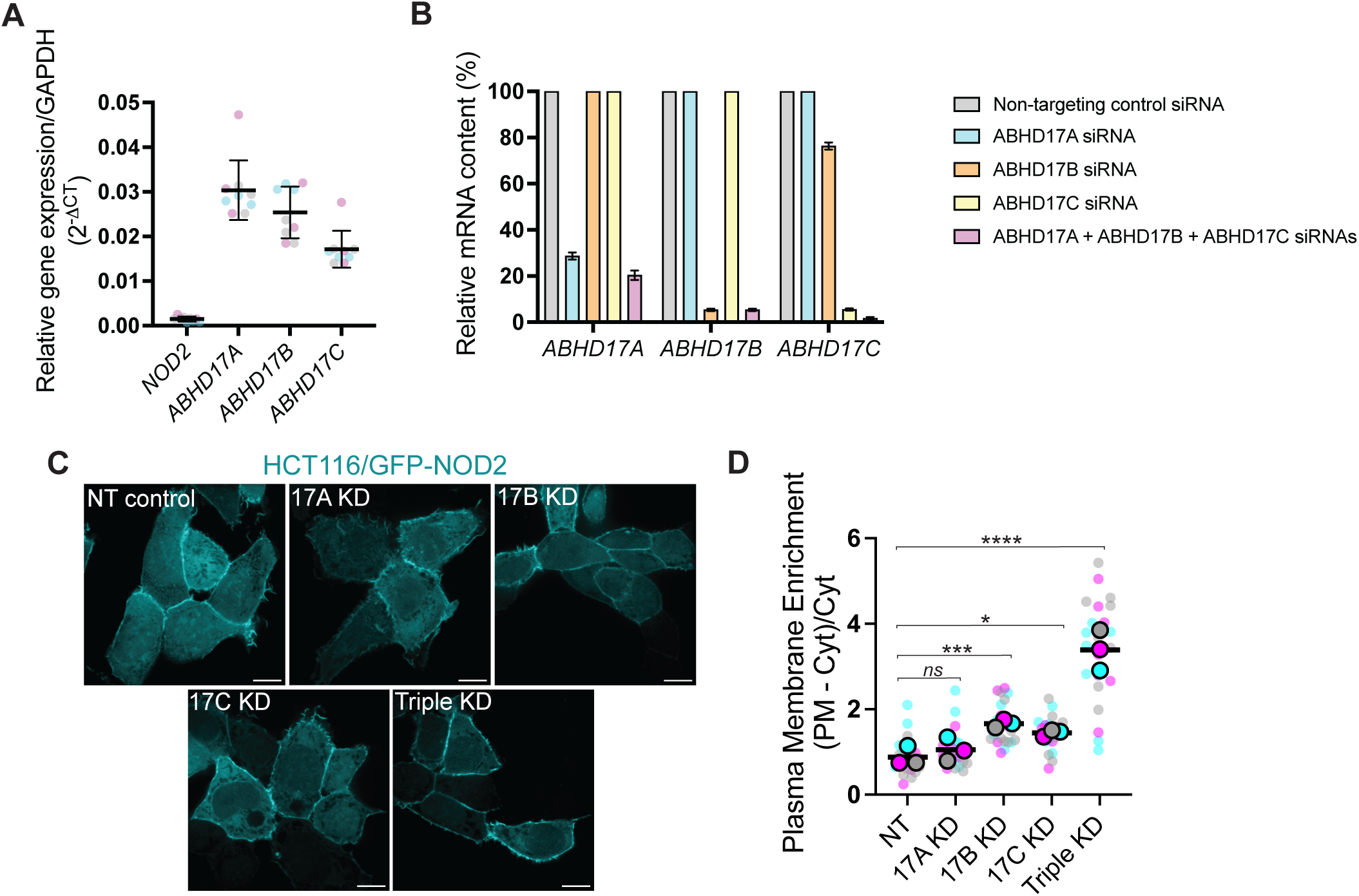
SiRNA-mediated ABHD17 knockdown enhances the plasmalemmal localization of GFP-NOD2 in HCT116 cells. (*A*) Quantitation of mRNA abundance for *NOD2, ABHD17A*, *ABHD17B* and *ABHD17C* gene expression relative to *GAPDH* (2^-ΔCT^) in HCT116 cells. (*B*) RT-qPCR of *ABHD17A*, *ABHD17B,* and *ABHD17C* transcript levels in HCT116 cells treated with small interfering RNA (siRNA): non-targeting (grey), ABHD17A (cyan), ABHD17B (orange), ABHD17 C(yellow) or a mixture of all three siRNAs targeting ABHD17A-C (magenta) for 48h. (*C*) Confocal images of GFP-NOD2 in HCT116 treated with the indicated siRNAs. Scale bars, 10μm. (*D*) Quantification of GFP- NOD2 plasma membrane localization from images as in *C*. All data are representative of at least three independent experiments. Mean values ± SD in *A*. p-values for data in *D* were determined by one-way ANOVA followed by Dunnett’s multiple comparisons tests. Group differences were considered significant for p≤0.05 (*), p≤0.001 (***) and p≤0.0001 (****). Not significant; *ns*.

### Attenuating ABHD17 Increases NOD2 Signaling

Next, we examined how inhibiting or silencing ABHD17 would affect endogenous NOD2-dependent activation and downstream pathways. To do this, we monitored the phosphorylation status of downstream effector proteins NF-κB (p65), IκBlil, and MAPK (p38) in response to NOD2 – RIPK2 activation (*Figure 5A-D*). Previous research has shown that inhibiting *S-*acylation of NOD2 reduces NF-κB and MAPK activity in RAW267.4 and HCT116 cells stimulated with MDP [21]. Next, HCT116 cells were treated with siRNAs or ABD957, as in Figure 4, before stimulation with a synthetic NOD2 agonist, L18-MDP. As predicted, cells with attenuated ABHD17 and elevated NOD2 *S*-acylation displayed an increase in the activation of NF-κB and MAPK responses (*Figure 5B-D*). Ultimately, this signal transduction cascade results in the upregulation of cytokines; therefore, we examined the secretion of cytokines under these conditions. The knockdown of individual isoforms was insufficient to promote increased interleukin 8 (IL-8), but did result in enhanced granulocyte-macrophage colony-stimulating factor (GM-CSF) production and secretion (*Figure 5E, F*). Silencing all three isoforms further enhanced the production and secretion of both IL-8 and GM-CSF. Again, to complement the siRNA-mediated knockdown experiments, we conducted an additional experiment using ABD957. Cells treated with ABD957 before ligand stimulation secreted ≈ 6-fold (p<0.0001) more IL-8 than DMSO-treated HCT116 cells (*Figure 5G*) and ≈ 3-fold (p<0.0001) more GM-CSF (*Figure 5H*). Together, these results demonstrate that the loss of ABHD17 function increases NF-κB and MAPK signaling pathways in response to the NOD2 ligand MDP.

**Figure 5.**
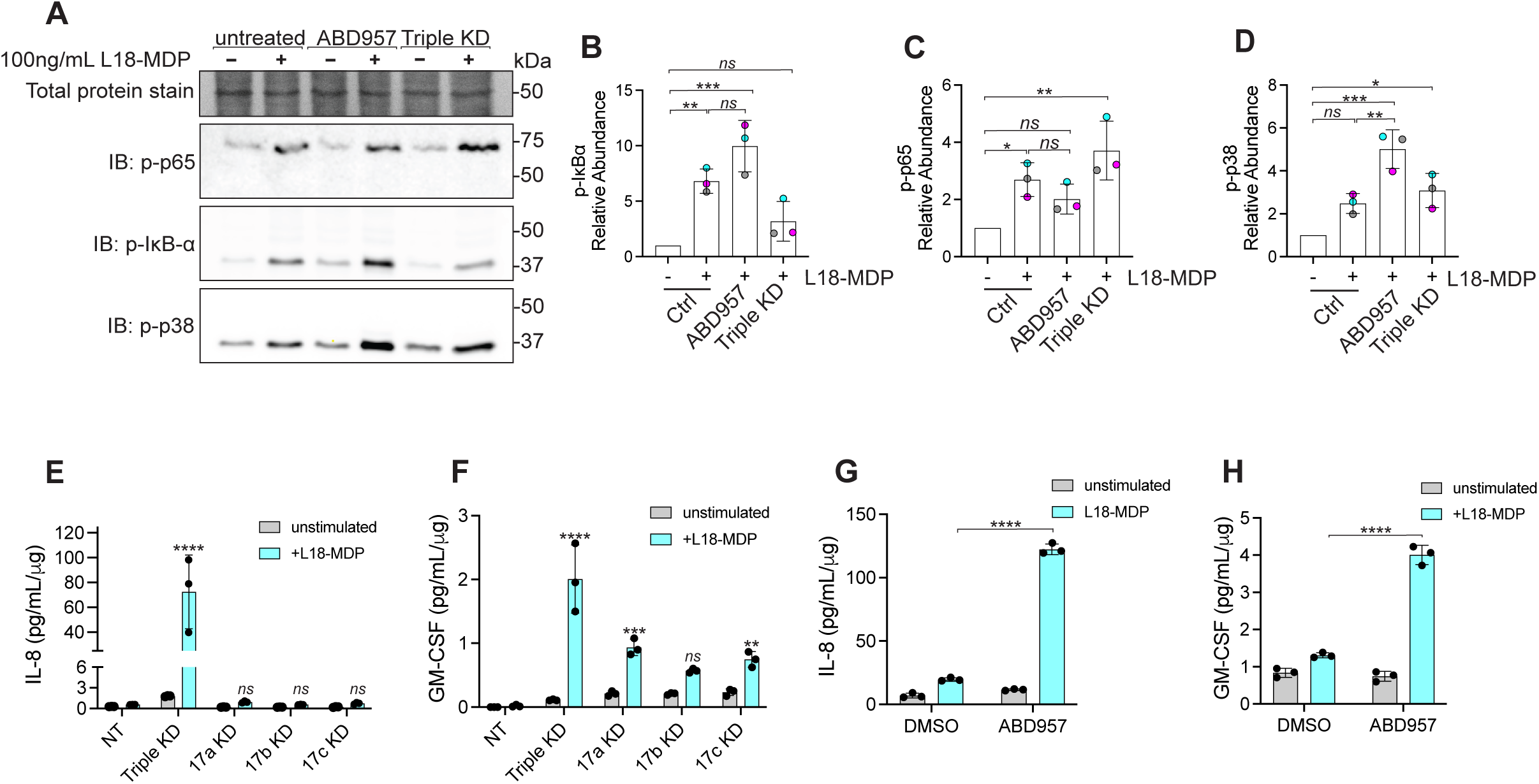
SiRNA-mediated ABHD17 knockdown enhances NOD2 signaling in HCT116 cells. (*A*) Immunoblots of phosphorylated- IkBα, NF-κB p65, and MAPK p38 in HCT116 cells. (B-D) Quantification of immunoblots as in A. (*E-H*) Cytokine (GM-CSF and IL-8) levels in cell culture supernatants. All data are representative of at least three independent experiments. All error bars, mean values ± SD. P-values for data in *B, C* & *D* were determined by one-way ANOVA followed by Tukey’s multiple comparisons test. Statistical significance of data in E, F, G & H was determined by two-way ANOVA then Šídák’s multiple comparisons test. Group differences were considered significant for p≤0.05 (*), p≤0.01 (**), p≤0.001 (***) and p≤0.0001 (****). Not significant; *ns*.

### Inhibiting ABHD17 Enzymes Restores Localization and S-acylation of select CD-Associated NOD2 Variants

Mechanistically, mutations in NOD2 could manifest in loss-of-function in numerous ways. For example, mutations could abrogate MDP binding, RIPK2 binding, ATP binding, or *S*-acylation. Indeed, it was previously reported that several CD-associated NOD2 loss-of-function variants have impaired S-acylation and do not localize to the plasma membrane [21]. Targeted inhibition of ABHD17 enzymes could increase the *S*-acylation of hypo-acylated mutants, potentially improving their ability to respond to MDP. To examine this possibility, we again used the Sleeping Beauty transposon system to create a stable transgenic cell line for each mutant using HCT116 and HEK293T cells. We examined several hypo-acylated variants with mutants spread throughout the protein, including the relatively common R702W and two other less common variants L248R, located between the CARD domains and the NACHT domains, and A755V, located between the NACHT and LRRs. We used confocal microscopy to examine the effects of inhibitors (THL, ABD957 and Palmostatin B) and RNA interference on the subcellular localization of each variant (*Figure 6A*). As we observed with wildtype NOD2, inhibiting ABHD17 increased the recruitment of L248R, R702W, and A755V to the plasma membrane (*Figure 6B-D*). Contrary to what we observed with wildtype NOD2, THL had no significant impact on the localization of the NOD2 mutants (p>0.05). We also confirmed that increased plasma membrane localization was associated with enhanced *S*-acylation (*Figure 6E-H*). These results suggest that some hypo-acylated NOD2 mutants are substrates for the acyltransferase ZDHHC5 and the ABHD17 thioesterase enzymes. Notably, inhibiting the deacylation mediated by the thioesterases increased their respective plasmalemmal localization.

**Figure 6.**
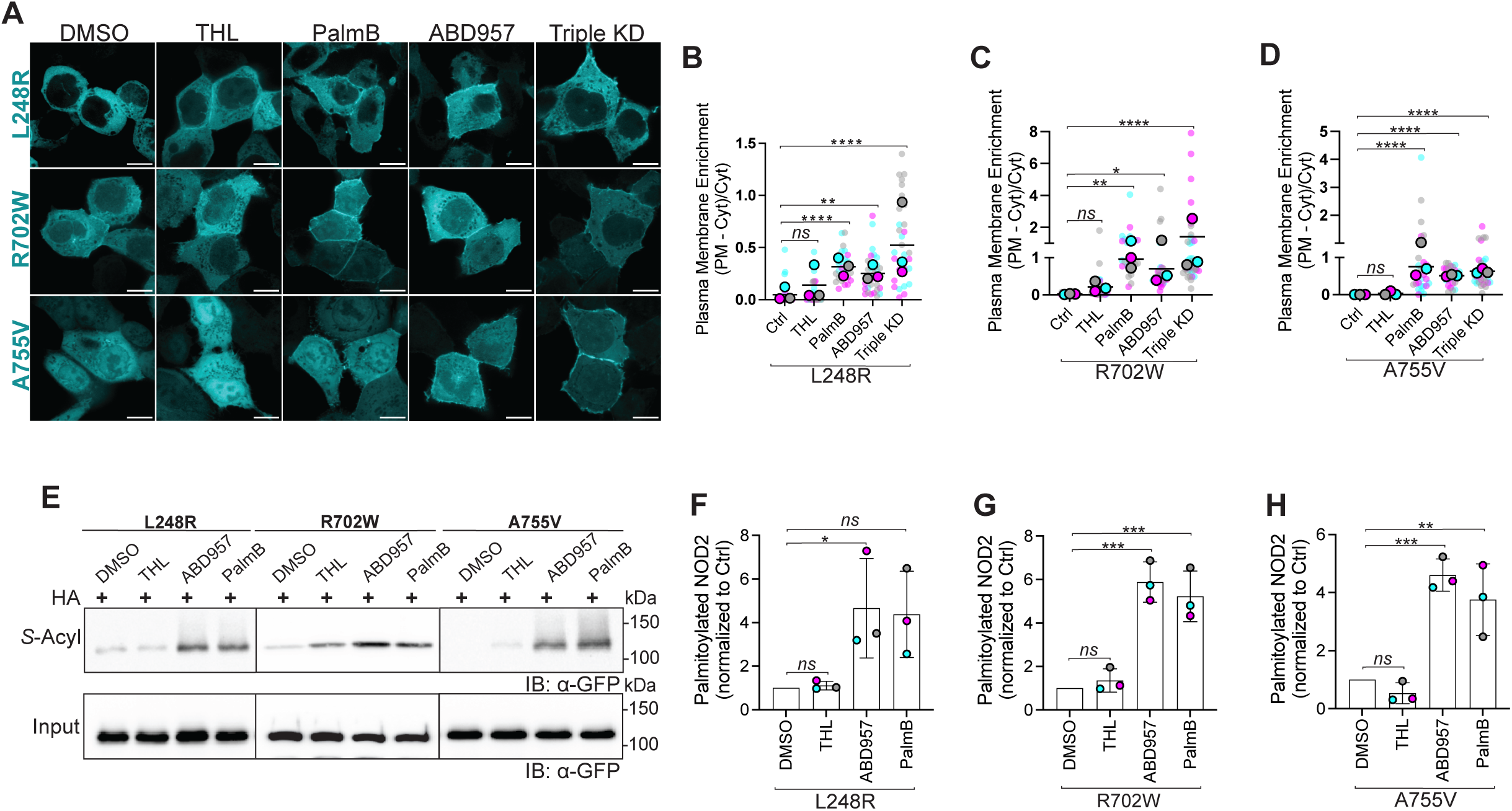
Inhibition of ABHD17 enzymes partly restores *S*-acylation of defective NOD2 variants. (*A*) Confocal images of doxycycline-inducible GFP-tagged human NOD2 variants L248R, R702W, and A755V in HCT116 cells. Cells were treated with solvent control, 40 μM THL, 100 μM PalmB, 2.5 μM ABD957, or a combination of siRNAs targeting ABHD17A, B, C (triple KD). Scale bars, 10 μm. (*B-D*) Quantification of plasma membrane localization of the GFP-NOD2 variants from images as in *A*. (*E-H*) *S*- acylation levels of NOD2 variants L248R, R702W and A755V in HCT116 cells treated with THL, ABD957 or PalmB. All data are representative of at least three independent experiments. Data are mean values ± SD. P-values for data in *B, C, D,* F, G & H were determined by one-way ANOVA followed by Dunnett’s multiple comparisons test. Group differences were considered significant for p≤0.05 (*), p≤0.01 (**), p≤0.001 (***) and p≤0.0001 (****). Not significant; *ns*.

Next, we sought to determine if increasing the targeting for these variants to the plasma membrane influences their ability to signal. For this, we used Sleeping Beauty- engineered stable doxycycline-inducible HEK293T cells, chosen for their minimal to non-existent endogenous NOD2 expression [7, 41], as a model for exploring agonist-induced NF-κB activation and cytokine production. Protein expression was induced with 10 ng/mL doxycycline for 18-20h. We titrated doxycycline to the lowest effective concentration to induce expression while preventing spurious activation of NOD2- mediated NF-κB signaling. Cells were stimulated with 100 ng/mL L18-MDP for 1h after RNA interference or a 4h treatment with ABD957. We show NOD2-RIPK2 activation of NF-κB and MAPK pathways, as in the HCT116 cells, using immunoblotting to capture phosphorylation of IκBlil and p38 (*Figure 7A-C*). Next, we analyzed the culture medium for pro-inflammatory mediators to correlate these acute stimulations with cytokine release from the HEK293T cells. IL-8 and TNFα were elevated compared with DMSO control (*Figure 7D*, *E*). HEK293T cells stably expressing A755V exhibited significant (p<0.0001) 4.1 and 3.4-fold increases in IL-8 and TNFα secretions when cells were stimulated with L18-MDP (*Figure 6D*, *E*). Cells expressing R702W displayed a modest increase in TNFα, and in IL-8 secretion. Cells expressing L248R also displayed a modest, albeit not statistically significant, in TNFα and IL-8 secretions when stimulated with L18-MDP. The results show that inhibiting these hypo-acylated NOD2 variants can partly increase their localization to the plasma membrane. However, the magnitude of signal transduction is variable at least in this engineered model.

**Figure 7.**
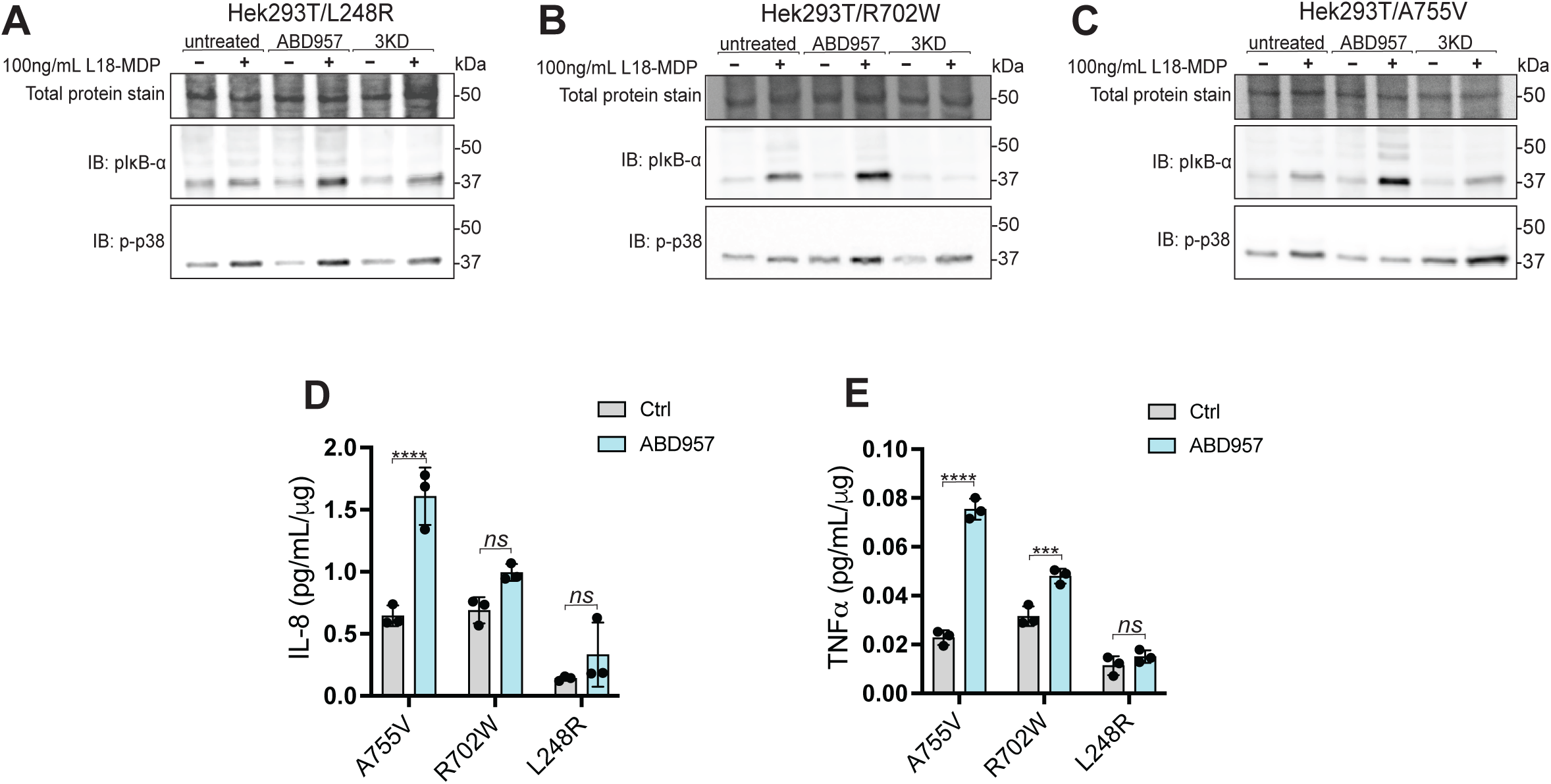
Targeting ABHD17 isoforms improves the functionality of select Crohn’s disease-associated NOD2 variants. (*A*) Detection of phosphorylated-IKB[] and phosphorylated-p38 MAPK using immunoblotting with the No Stain total protein stain. Whole-cell lysates of HEK293T cells stably expressing (*A*) L248R (*B*) R702W (*C*) A755V were simulated with L18-MDP for 1h after being treated with either DMSO, ABD957, or siRNAs. Cytokine levels (*D*) IL-8 and (*E*) TNFα in cell culture supernatants. All data and immunoblots are representative of at least three independent experiments. Data represents the mean ± SD. Significance of *D* & *E* was assessed by calculating P-values of the using two-way ANOVA then Šídák’s multiple comparisons test. Group differences were considered significant for p≤0.0001 (****). Not significant; *ns*.

### S-acylation deficient NOD2^L1007fs^ and NACHT domain mutants are not improved by ABHD17 inhibition

To better understand the acylation–deacylation cycle and how it may be altered in the different mutants, we next examined the CD-associated 3020insC frameshift mutation that encodes a C-terminally truncated NOD2 protein lacking the last 33 amino acids (L1007fs). This C-terminal truncation mutant lacks Cys1033, one of the two S-acylation sites in human NOD2. Previous reports demonstrate that this mutant is predominantly cytosolic and that the Cys395 is minimally acylated in HEK293 cells [21]. However, it is unknown if this is because this mutant is a poor substrate or if it is more rapidly deacylated. Examining stable doxycycline-inducible GFP-NOD2^L1007fs^ in HCT116 cells reveals no plasma membrane localization and is unresponsive to ABHD17 inhibition (*Figure 8A*). Consistent with the microscopy data, using acyl-RAC, we detect no *S*-acylated protein in the presence or absence of ABD957 (*Figure 8B-C*); although this maneuver increased both wild-type and A755V S-acylation levels. These results suggest that GFP-NOD2^L1007fs^ is likely a poor substrate for ZDHHC5 and raises the possibility that Cys395 is acylated downstream of Cys1033 acylation.

**Figure 8.**
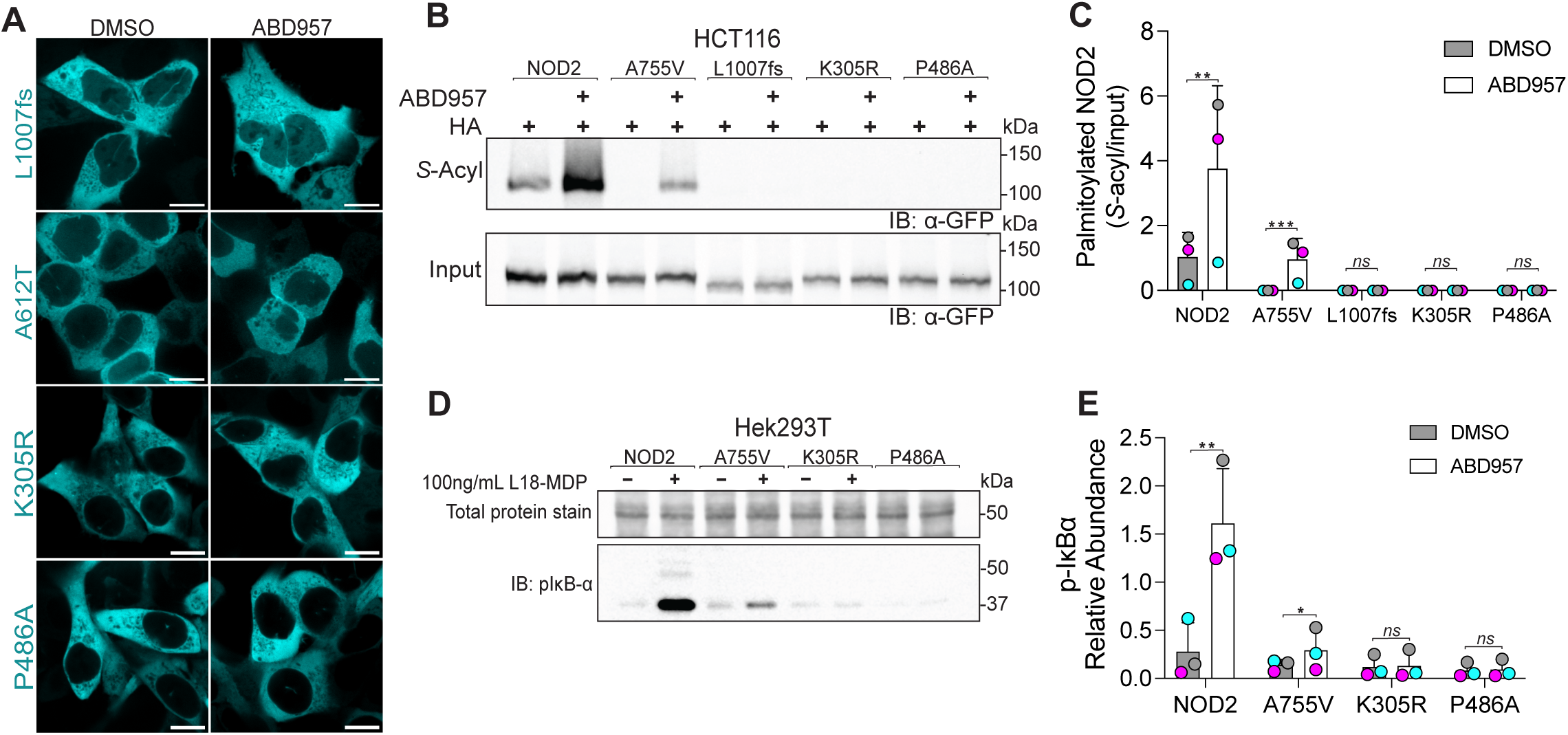
NOD2^L1007fs^ and NACHT mutants are not *S*-acylated or improved by targeting ABHD17 enzymes in HCT116 cells. (*A*) Confocal images of HCT116 cells stably transfected with doxycycline-inducible NOD2 variants: L1007fs and three NACHT/nucleoside-triphosphatase domain mutants, the human variant A612T, and two experimentally generated mutants K305R and P486A treated with DMSO or ABD957 for 4 hours. (*B*) S-acylation of wild-type NOD2 and the indicated mutants treated with ABD957 inhibitor using acyl-RAC and immunoblotting. Note: wild-type and A755V are used as positive controls in this experiment. (*C*) Quantification of panel *B* from three independent experiments. (*D*) HEK293T cells transiently expressing wt-NOD2 and the indicated variants were incubated with ABD957 +/- MDP for 1 h. Whole cell lysates were resolved and phosphorylated-IKB[] was detected by immunoblotting. All data are representative of at least three independent experiments and samples from individual replicates color-matched. An unpaired two-tailed Student’s t-test was used to compare the solvent control and ABD957 conditions for wt-NOD2 and each variant in *C* & *E*; Group differences were considered significant for p≤0.05 (*), p≤0.01 (**) and p≤0.0001. Not significant; *ns*.

Next, we examined three mutants residing within the central nucleotide-binding and oligomerization domain, also called the NACHT domain. Specifically, we examined the human variant A612V and two experimentally generated mutants, K305R and P486A. The K305R (Walker-A motif), P486A (GxP motif), and A612T (Walker-B motif) mutations span the NACHT domain and are required for ATP binding and hydrolysis [42]. Using the Sleeping-Beauty doxycycline-inducible system, we examined the localization of GFP chimeras of these three mutants. Like the GFP-NOD2^L1007fs^, these mutants display no plasma membrane localization in resting cells, which does not change after 4 hours of treatment with ABD957 (*Figure 8A*). Using acyl-RAC, these NACHT domain mutants show no basal *S*-acylation and no increase following incubation with ABD957, in contrast to wild-type or the A755V variant (*Figure 8B, C* and 6*E, H*). Consistent with previous reports [42], the K305R and P486A failed to activate the NF-κB pathway, and treatment with ABD957 could not increase signaling (*Figure 8D, E*). These results are consistent with the need for an initial ATP-binding and/or hydrolysis for NOD2 *S*-acylation.

## DISCUSSION

The *S*-acylation of NOD2 attaches the protein to the plasma membrane, endosomes, and phagosomes critical for MDP-induced signaling [21]. Here, we demonstrate that ABHD17 enzymes remove the two acyl chains from NOD2, thus negatively regulating NOD2-mediated NF-lrlB signaling. The results further highlight how the S–acylation–deacylation cycle is a pivotal regulator of NOD2 function. Specific poorly *S*-acylated variants can be improved by targeting the ABHD17 family of thioesterases. This study further supports the previous findings that defective NOD2 *S*- acylation renders the protein non-functional in response to MDP and may contribute to CD’s pathogenesis. Our findings also raise the possibility that approaches that attenuate the deacylation of NOD2 variants may be beneficial therapeutically depending on the specific NOD2 mutation. It is worth noting that our work has been conducted in immortalized intestinal epithelial cells and HEK cells. This limitation could be mitigated by generating new mouse models or using CRISPR to generate iPSC cells with specific NOD2 variants for testing in organoid models.

This study investigated NOD2 loss-of-function variants associated with CD, including two more prevalent NOD2 mutations, L1007fs and R702W. The C-terminal truncation of 1007fs deletes one of the two *S*-acylation sites. Both these mutant proteins, in addition to L248R and A755V, are hypo-acylated and mislocalized in cells and have lost the ability to induce NF-lrlB signaling in response to MDP [21]. Although the previous study revealed some residual *S*-acylation of the FLAG-NOD2^L1007fs^ in Hela cells, we did not detect any in our system using the acyl-RAC assay. Furthermore, S-acylation was not increased in this mutant upon inhibition of ABHD17 isoforms. A few differences between the previous and current studies could result in differences. This includes different assays to assess *S*-acylation, different cell lines, and transient vs. stable inducible expression of the NOD2 proteins. However, the collective results demonstrate little to no *S*-acylation of the NOD2^L1007fs^ protein. Mechanistically, we can envisage a few possible explanations. First, the truncated protein is not recognized as a substrate by the ZDHHC5 enzyme. Second, there is a hierarchy in the order of *S*- acylation events, with Cys1033 being modified first and required for Cys395 to be acylated. Third, although our results suggest that ABHD17s are the primary thioesterases for NOD2, it is possible that a mono-*S-*acylated NOD2^L1007fs^ could be deacylated by another enzyme that usually doesn’t recognize the dually *S*-acylated NOD2 as a substrate. Regardless of the precise mechanism, it appears that the functionality of this mutant cannot be restored.

We observed improved *S*-acylation and plasmalemmal localization, and functionality in three hypo-acylated mutants, R702W, L248R and A755V. We also consistently found that the enhanced S-acylation also increase the phospho-IκB-α and a tendency to increase the secretion of cytokines in the HEK cells. In contrast, mutations within the NACHT domain display no *S*-acylation, which did not change with the inhibition of ABHD17 isoforms. It remains unknown which among the sixty CD-associated NOD2 variants are hypo-acylated and which could be improved through attenuation of ABHD17. Based on our findings, it is tempting to speculate that ZDHHC5 specifically recognizes and *S*-acylates the ATP-bound NOD2. Ultimately, we suspect that targeted inhibition of ABHD17 could benefit a subset of these human variants.

The connection between NOD2 loss-of-function mutants and CD may have significant implications for the disease. This is not only because dysfunction in the innate immune system is crucial in CD’s development but also because it has been demonstrated that restoring normal innate signaling can alleviate colitis [43]. However, ongoing debates exist regarding how changes in NOD2 signaling can increase susceptibility to inflammation and the role post-transcriptional regulation plays in patients with or without NOD2 variants associated with the disease and their resulting pathogenic innate immune responses [44, 45]. *S*-acylation has been shown to impede autophagic degradation of NOD2 by attenuating binding to the cargo receptor p62/SQSTM1 [46]. The degradation rate is higher for *S*-acylation-deficient NOD2 variants than wild-type NOD2 [46, 47]. Thus, ABHD17 enzymes may also regulate NOD2 stability by coupling deacylation to its autophagic removal.

Additional roles for NOD2 have been described in response to viruses and ER stress. These roles for NOD2 do not require localization to the PM or endocytic organelles; instead, NOD2 redistributes to the mitochondria and ER. It is unclear whether these responses require *S*-acylated NOD2 or if ABHD17 isoforms will influence these pathways. Teleologically, both plasmalemmal (*S*-acylated) and cytosolic (non-acylated) pools of NOD2 may exist to allow NOD2 to function in many subcellular compartments under various stimuli and conditions. However, this requires further experimentation to assess this possibility.

## ACKNOWLEDGEMENTS

The authors thank the Kennan Research Center for Biomedical Science Core Facilities at St. Michael’s Hospital, especially Dr. Caterina di Ciano-Oliveira and Dr. Monika Lodyga, for their continued support, technical advice, expertise, and training.

## AUTHOR CONTRIBUTIONS

**Charneal Dixon** investigation, formal analysis, manuscript first draft. **Noah Martin** investigation. **Micah Niphakis** resources. **Benjamin Cravatt** resources and funding acquisition. **Gregory Fairn** conceptualization, writing–review & editing, supervision, project administration, and funding acquisition.

## RESOURCE AVAILABILITY

### Lead contact

Further information and requests for resources and reagents should be directed to and will be fulfilled by the lead contact, Gregory Fairn

### Materials availability

- Any plasmids or cell lines generated in this study are available upon request.

### Data and code availability

- This paper does not report any original code.
- Any additional information required to reanalyze the data reported in this paper is available from the lead contact upon request.

## METHODS

### Reagents

L18-MDP (cat# tlrl-lmdp) was purchased from Invivogen (San Diego, USA). ML348 (cat# 899713-86-1), ML349 (cat# 890819-86-0), WWL70 (cat# 947669-91-2), orlistat (cat# 96829-58-2), *S*-Methyl thiomethanesulfonate (cat# 64306), sodium dodecyl sulfate (cat# 71736-500), hydroxylamine (cat# 159417), 4’,6-diamidine-2’-phenylindole dihydrochloride (cat# 10236276001), NP-40 (cat# 492016-500L), Triton X-100 (cat# X- 100) and puromycin (cat# P4512) were purchased from Sigma-Aldrich (Oakville, Canada). Palmostatin B (cat# 178501) and RIPA (cat# 20-188), products of EMD Millipore, were also purchased through Sigma-Aldrich (Oakville, Canada). Doxycycline hydrochloride (cat# DB0889) was obtained from BioBasic Inc. (Markham, Canada).

High-capacity acyl-RAC S3 capture beads (cat# AR-S3-1) were purchased from NANOCS (Natick, USA).

### Plasmids

ABHD17 (17A, 17B & 17C), ABHD6, ABHD12 and ABHD16A in pEF-BOS-HA were gifts of Dr. Jennifer Greaves (Research Centre for Health and Life Sciences, Coventry University, Coventry, U.K.). These plasmids encode full-length human ABHD17 (17A, 17B & 17C), ABHD6, ABHD12 and mouse ABHD16A. pSB-GFP-NOD2 was generated by cloning GFP-NOD2 into pSBTet-BP using two SfiI restriction sites on either side of the luciferase cassette. The NOD2 mutants were generated in this vector using the Q5 site-directed mutagenesis kit from New England Biolabs, resulting in single amino acid changes L248R, K305R, P486A, R702W, A612T, A755V and 3020insC in the expressed proteins.

### Immortalized and engineered cell lines

Cells were purchased from the American Type Culture Collection (ATCC).

Human colorectal carcinoma HCT116, HT29 and human embryonic kidney (HEK) 293T cell lines were maintained and propagated in high glucose Dulbecco’s modification of Eagle’s medium (DMEM) from Wisent (St. Bruno, Canada, cat# 319-020-CL) supplemented with 1.5 g/L sodium bicarbonate, L-glutamine and 10% fetal bovine serum (Wisent, cat# 080150) in a humidified incubator at 37 °C, 5% CO_2_.

Stable cells were generated using the sleeping beauty transposon system [48].To generate doxycycline-inducible stable integration of GFP-NOD2, cells were co-transfected with sleeping beauty vector (pSBTet-BP [48], Addgene # 60496) carrying the gene of interest and small amounts of the transposase encoding plasmid, pCMV(CAT)T7-SB100 [49] (Addgene # 34879). According to the manufacturer’s instructions, cells were transfected using ViaFect transfection reagent (Promega, Madison, WI). Forty-eight hours after transfection, cells were selected for puromycin resistance over 3-5 days and terminated when virtually all the cells displayed the expected doxycycline inducible GFP or constitutive BFP fluorescence. Expression of the GFP-NOD2 and the indicated mutants was induced using 1 μg/mL doxycycline unless otherwise indicated. Stable expression of GFP chimeric proteins was confirmed using microscopy and immunoblotting.

### Transfections

cDNA transfections were performed using Promega ViaFect (cat# E4982), while siRNA transfections were performed using DharmaFECT2 (cat# T-2002-02) purchased from Horizon Discovery Biosciences. HCT116 cells were transfected with cDNAs as indicated in each experiment using ViaFect as per manufacturer’s instructions. For immunofluorescence studies, HCT116 cells were grown on glass coverslips in 12-well plates and transfected at 60–80% confluence with 1 µg of cDNA per well for each of the indicated thioesterase encoding plasmids using Promega’s Viafect according to product instructions.

Unless otherwise stated, for RNA interference experiments two rounds of transfection were used to introduce 100nM small interfering RNA in HCT116 and HEK293T on days 1 and 2 post-plating. For knockdown experiments, the ON-TARGETplus siRNA of targeted genes and the non-targeting siRNA were purchased from Horizon Discovery Biosciences. The silencer siRNA used are as follows: ON-TARGETplus Non-targeting siRNA, 5’-UGGUUUACAUGUCGACUAA- 3’, ON-TARGETplus Human ABHD17A siRNA, 5’-ACGUGAAAACGGAAAUUAA-3’, ON- TARGETplus Human ABHD17C siRNA, 5’-CCACCAAUGUUCUCGGUAU-3’, ON-TARGETplus SMARTpool Human ABHD17B siRNA (1# 5’- GGACUAGGAUCACGGAUUA-3’, 2# 5’-GGAUCUUGCUGCUCGAUAU-3’, 3# 5’-GUUCACCCAAUGCGAAAUA-3’, 4# 5’-UCGCAUUGUUUGAACGUUG-3’). Since isoform-specific antibodies against ABHD17 isoforms are not available, RT-qPCR was used to quantify the degree of mRNA knockdown.

Experimentally, cells transfected with siRNA were induced with doxycycline on day 3, followed by confocal microscopy, cells were stimulated with L18-MDP on day 4, 18-20h following induction.

### Inhibitor assays

Stable transfected HCT116 were cultured as monolayers on 18mm round glass coverslips in 12-well plates. Stable expression of GFP-tagged NOD2 or NOD2 variant was induced with doxycycline (1 μg/mL). Approximately 18-20h after induction, cells were treated with DMSO vehicle (control) or small molecule inhibitors Palmostatin B, Orlistat (tetrahydrolipstatin; THL), ABD957, WWL70, ML348, ML349 for 4 h at 37°C in 5% CO_2_.

### Preparation of cell extracts

Stable transfected HCT116 and HEK293T cells were induced with doxycycline (1 μg/mL) for 18-20 h, then treated with inhibitors or stimulated as indicated. Cells were harvested in phosphate-buffered saline (D-PBS; cat# 311-010-CL) from Wisent Inc., and lysed in RIPA buffer (CST; cat# 9806S) or NP-40 lysis buffer containing PhosSTOP phosphatase inhibitor (Roche; cat# 04906845001) and EDTA-free protease inhibitor (BioShop; cat# PIC002-1) and incubated on ice for 1 hour, then centrifuged at 10,000 x*g* at 4°C for 10 minutes to obtain clear lysates. Protein concentration was determined with Pierce BCA protein assay (Thermo Scientific; cat# 23225).

### Confocal microscopy

Cells were seeded on 18 mm-round glass coverslips in 12-well plates. Stable HCT116 cells expressing GFP-NOD2 wildtype and mutants were washed with PBS and fixed with a 4% paraformaldehyde solution for 20 min at room temperature. Glass coverslips were washed with PBS then incubated with 100 mM glycine, 0.1% v/v Triton X-100 for 20 min at room temperature. Cells were blocked with 5% BSA in PBS for 1h at room temperature and then probed with 1:100 anti-HA (CST) antibody overnight. The coverslips were washed three times with PBS, followed by a 1h incubation with species-matched 1:500 Cy5-conjugated goat anti-mouse IgG (Jackson ImmunoResearch) in PBS. Images were acquired in the St. Michael’s Hospital Bioimaging Facility using a Zeiss Axiovert 200M microscope equipped with a Yokogawa CSU-X1 confocal spinning disc system and a 63X Apo PLAN VC, NA 1.40 oil objective and a Hamamatsu ImagEM X2 EM-CCD camera. Images were processed using ImageJ2 [50] version 2.14.0/1.54f.

Acyl resin-assisted-capture (Acyl-RAC) Stable HCT116 cells were harvested and lysed (1% Triton X-100, 50 mM Tris- HCl (pH 7.5), 150 mM NaCl, 1 mM EDTA, 1 % SDS, 0.5 % sodium deoxycholate, 1X protease inhibitor (BioShop; cat# PIC002-1), 0.2% v/v methyl methanethiosulfonate (MMTS). Lysates were sonicated and incubated at 42°C in a thermal shaker for 3h. Blocking was stopped, and MMTS was removed with 3-4 rounds of methanol/chloroform precipitation. The protein pellets were rinsed with methanol and air-dried. The pellets were redissolved in binding buffer (100 mM Tris- HCl (pH 7.5), 1 mM EDTA, 1 % SDS) by incubating at 42°C in a thermal shaker, with intermittent mixing. The samples were sonicated in a water bath for 5-10 min to resuspend any undissolved pellet further. Each sample was divided in two, and 50 μL of hydroxylamine (NH2OH, 400 mM final concentration; or NaCl for control samples) was added before incubation with 25 μL of packed acyl-RAC beads (High-Capacity Acyl-RAC S3 beads; Nanocs # AR-S3-1,2) for 2 h at room temperature. The beads were washed 3 times with binding buffer before adding 4X SDS sample buffer with DTT and incubating the slurry at 80°C for 15 minutes.

### RNA extraction and RT-qPCR

Total RNA was extracted from HCT116 cells RNeasy Mini Kit (cat# 74104; Qiagen, Germany) according to the manufacturer’s protocols. cDNA was synthesized from 1 µg RNA using SensiFAST cDNA Synthesis (cat# BIO-65053; FroggaBio; Concord, Canada) and SensiFAST SYBR No-ROX kit (cat# BIO-98005). qPCR was performed using a BIO-RAD CFXOpus 96 Real-Time PCR System in conjunction with CFX Maestro software version 5.0.0.21.0616 (BIO-RAD; Ontario, Canada). All cycle threshold (CT) values were normalized to GAPDH. The relative expression level for each gene of interest was calculated using the 2^-ΔΔC^T method [51]. The primer sequences were as follows: Human *ABHD17A* forward: 5’- ACCATCGAGGTCTTCCCCACCAGG-3’, Human *ABHD17A* reverse: 5’- AGAAGAGGACCGTGTACCTGGCACC-3’, Human *ABHD17B* forward: 5’- CCGCTCAGGTGCAGAGCATGAATAA-3’, Human *ABHD17B* reverse: 5’- AAGCAATCTTCCCTGGACAAGGTGG-3’, Human *ABHD17C* forward: 5’- TTGATGCTTTCCCCAGCATTGACA-3’, Human *ABHD17C* reverse: 5’- TCGCTAGGCCATGGGAGAAATCG-3’, Human *GAPDH* forward: 5’-CAATGACCCCTTCATTGACC-3’, Human *GAPDH* reverse: 5’- GACAAGCTTCCCGTTCTCAG-3’, Human *NOD2* forward: 5’- GCACTGATGCTGGCAAAGAACG-3’, Human *NOD2* reverse: 5’- CTTCAGTCCTTCTGCGAGAGAAC-3’.

### Multiplex analysis of secreted cytokines

Using 12 well tissue culture plates, cells were cultured with DMEM containing 10% FBS. Cells were treated with siRNA complexes over 48 hours or ABD957 inhibitor for 4 hours, before brief stimulation with 100 ng/mL L18-MDP (1h for HEK293T cells and 6h for HCT116 cells). Plates were maintained at 37 °C in 5% CO_2_ for 24 h. Next, supernatants were collected and analyzed using Luminex-100 MAP^®^ technology for a panel of 14 human cytokines, chemokines, and growth factors. Specifically, plates were placed on ice, the contents of the wells for each condition (unstimulated, L18-MDP- stimulated) were transferred to 1.5mL microcentrifuge tubes, and samples were centrifuged at 4 °C for 10 min at 11,000 × *g* followed by distribution of the supernatant into 200 µl aliquots that were immediately frozen at −80 °C and maintained frozen until analysis. This was repeated 3 times to obtain 3 independent replicates. This multiplexing analysis was performed using a Luminex™ 200 system (Luminex, Austin, TX, USA) by Eve Technologies Corp. (Calgary, Alberta). Fourteen markers were simultaneously measured in the samples using Eve Technologies’ Human High Sensitivity 14-Plex Discovery Assay® (MilliporeSigma, Burlington, Massachusetts, USA). The 14-plex consisted of GM-CSF, IFNγ, IL-1β, IL-2, IL-4, IL-5, IL-6, IL-8, IL-10,IL-12p70, IL-13, IL-17A, IL-23, TNF-α. Assay sensitivities of these markers range from 0.11 – 3.25 pg/mL for the 14-plex, and some samples required dilution for accurate determination. Individual analyte sensitivity values are available in the MilliporeSigma MILLIPLEX® MAP protocol. Only molecules that reached the low end of the sensitivity are depicted in the results.

## QUANTIFICATION AND STATISTICAL ANALYSIS

Statistical differences (p<0.05) were inferred using Student’s two-tailed t-tests or one-way ANOVA with Dunnett’s correction or two-way ANOVA with Fisher’s least significant difference (LSD) method for multiple comparisons performed using Prism 10 (Graph-Pad Software, Inc., La Jolla, CA) with DMSO-treated, vector-co-transfected, or non-targeting siRNA-transfected samples as the control group. All significant differences are indicated in the text and figures. The statistical significance threshold was set to 0.05 throughout this study.

Cell membrane fluorescence intensity and immunoblot band intensities were quantified using ImageJ2 open-source image processing software.

