## Supplemental Figure 1 for "Attenuating ABHD17 isoforms augments the *S*-acylation and function of NOD2 and a subset of Crohn’s disease-associated NOD2 variants"

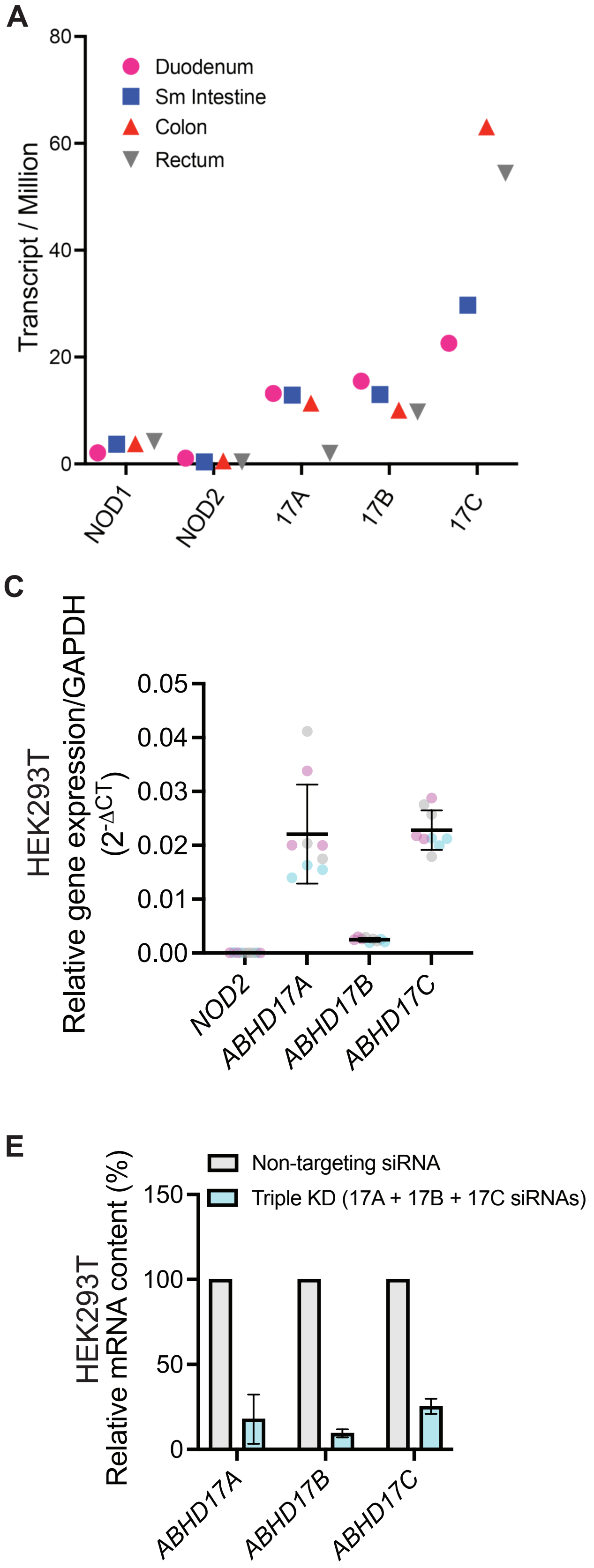


**Figure S1**. (*A, B*) Consensus RNASeq data for select tissues available from Human Protein Atlas (proteinatlas.org). (*C, D*) Quantitation of mRNA abundance for NOD2, ABHD17A, ABHD17B and ABHD17C gene expression relative to GAPDH (2-ΔCT) in HEK293T and HT29 cells. (*E, F*) RT-qPCR of ABHD17A, ABHD17B, and ABHD17C transcript levels in HEK293T and HT29 cells treated with a mixture of all three small interfering RNA (siRNA) targeting ABHD17A-C (cyan) for 48h.
